# Fine-scale recombination rates inferred using the canFam4 assembly are strongly correlated with previous maps of dog recombination

**DOI:** 10.1101/2025.09.11.675639

**Authors:** Jeffrey M. Kidd

## Abstract

**Background:** Genetic maps are a key resource for the analysis of genetic variation. Prior maps in dogs were based on the canFam3.1 genome assembly, which contains thousands of sequence gaps that include functional elements. Although long-read technologies have enabled the generation of updated reference assemblies, updated maps are required to take full advantage of these new resources.

**Methods:** We inferred fine-scale recombination maps that consider population demographic history for four populations of village dogs sequenced by the Dog10K Consortium. We compare properties among the newly generated and previously published maps and assess the effects of genetic map choice on genotype phasing accuracy.

**Results:** The newly inferred maps are broadly consistent with previous studies of recombination in dogs. At small scales, stronger correlations are observed among maps inferred using linkage disequilibrium than with a pedigree-based map. The position of recombination hotspots showed significant overlap among inferred maps and were enriched for sequences that were gaps in a previous canine genome assembly. When using the Dog10K samples as a reference panel, a phase error rate of approximately 2% was found with the choice of genetic map having little effect.

**Conclusions:** We describe genetic maps based on the canFam4 assembly which will be useful for future studies of genetic variation that rely on updated dog genome references.

## 1. Introduction

Recombination is essential for the proper segregation of chromosomes during meiosis and for the generation of genetic variability. The rate of recombination along the genome (i.e., cM per Mb) is known to vary widely between species, between males and females, and between individuals [1,2]. Genetic maps, which link physical distance along a chromosome with the frequency of genetic recombination, are a key resource for trait mapping and evolutionary studies.

Over the past two decades, domestic dogs have emerged as a powerful system for linking genotypes to phenotypes and for understanding the effects of selection and population history on genome variation [3-5]. This has catalyzed the development of key canine genomic resources including a well-annotated reference genome [6,7], detailed phenotype information coupled with databases of known genetic variation that enable efficient genome wide trait mapping [8], and publicly available collections of samples with short-read whole genome sequence data [9-11]. This strong research foundation has been expanded by the availability of multiple high-quality reference genomes derived from long-read sequencing technologies [12-22], consistently processed short-read sequencing data from a large collection of dogs and wolves [11], and ancient DNA sequences obtained from globally distributed samples of dogs and wolves [23-30].

Genetic maps are a key resource for the analysis of genetic variation. The most common approaches to estimate a genetic map include tracking recombination through pedigrees or analyzing patterns of linkage disequilibrium (LD) among markers genotyped in a population. Pedigree maps involve fewer assumptions than maps based on LD and can measure sex-specific recombination rates. However, the resolution of pedigree maps is limited by the number of recombination events that occur in the sample. Maps based on LD consider all recombination events that have occurred in the evolutionary history of the analyzed samples, permitting the estimation of recombination rates at fine scales that are averaged between sexes. Importantly, LD-based methods indirectly measure the absolute recombination rate, instead estimating a population recombination parameter that depends on the effective population size and can be biased by selection, mutation, and changes in demographic history [1,2].

In many eukaryotes recombination occurs in narrow physical regions known as hotspots [31]. The *Prdm9* zinc-finger protein is a major factor specifying the location of recombination hotspots in many metazoans, including humans [32-34]. However, the dog genome does not contain a functional copy of *Prdm9* [35]. Prior analyses have shown that although *Prdm9* was lost early in canid evolution recombination in dogs is concentrated into specific locations. These hotspots appear to be stable over long time scales and are characterized by high GC content and reduced DNA methylation [36,37]. Similar patterns have been found in other species that lack functional *Prdm9* alleles [38-40].

In 2010, Wong et al. constructed a canine genetic map based on ∼3,000 microsatellites and a mean of 266 informative meioses [41]. The resulting map enabled subsequent linkage studies that mapped the genetic basis underling canine traits. This map described a pattern of a low recombination rate at the centromeric end and a high rate of recombination at the telomeric end of canine autosomes and found that GC sequence content was correlated with recombination rate.

A widely used fine-scale map of recombination in dogs was created by Auton et al. in 2013. This map inferred recombination rates from patterns of linkage disequilibrium between SNPs found in 51 village dogs using whole genome sequencing [42]. Inference was performed using the *LDhat* software package [43,44], with all 51 samples analyzed together. The resulting map was broadly consistent with prior maps and confirmed a strong association between recombination rate and GC content, particularly for CpG islands near gene promoters. Auton and colleagues also described a skew in the rate of AT vs GC variation near recombination hotspots, consistent with the action of biased gene conversion in shaping the sequence composition of the canine genome. Despite the presence of hotspots located at regions of high CG content, the Auton et al. map suggested that recombination in dogs may be more uniform than recombination in humans. However, simulations showed that inferences using *LDhat* are sensitive to the effective population size of the studied population, with inferences from populations with larger effective sizes yielding more uniform maps.

The relative uniformity of recombination in dogs and humans was further explored in 2016 by Campbell et al. The Campbell genetic map was created using SNP genotype data from an extended dog pedigree with 408 informative meioses [45]. The Campbell pedigree map is more skewed than the Auton LD map, and comparison with a similarly constructed human pedigree map shows that recombination is less uniform in dogs than in humans.

Both the Auton LD and Campbell pedigree maps were constructed using the canFam3.1 genome assembly, derived from a Boxer breed dog [6,7]. Although a key enabling resource for the canine genomics community, this assembly has over 19,000 gaps, including in gene promoter regions. Analysis using long-read sequencing technologies showed that these gaps contain sequence with a high GC content and are enriched near recombination hotspots inferred by the Auton LD map [13]. Multiple high-quality dog and wolf assemblies have been described in recent years, with several large-scale sequencing projects relying on the UU_Cfam_GSD_1.0/canFam4 assembly derived from a German Shepherd Dog [14].

Although *LDhat* has enabled the inference of fine scale genetic maps in multiple species, it has several limitations including computational run time and the assumption of a constant effective population size. In this study we use *pyrho* [46], an efficient method for inferring recombination rates from LD patterns that accounts for changes in effective population size, to infer a new canine genetic map based on the canFam4 assembly. We apply this approach to four populations of village dogs sequenced by the Dog10K Consortium and compare the properties of the newly inferred recombination maps with versions of the Auton LD and Campbell pedigree maps converted to the canFam4 assembly.

## 2. Materials and Methods

### 2.1 Generating LD-based genetic maps for the canFam4/UU_Cfam_GSD_1.0 assembly

Canine genetic maps were inferred from patterns of linkage disequilibrium between SNPs using *pyrho* (v0.2.0) [46,47]. SNP genotypes were generated by the Dog10K Consortium based on alignment to the UU_Cfam_GSD_1.0 (canFam4) genome assembly derived from a German Shepherd Dog [11]. SNPs that overlap with regions identified as segmental duplications were removed [48]. Maps were made separately for four populations of village dogs (Iran, n=24 samples; Kenya, n=18 samples; Liberia, n=17 samples; and Congo, n=16 samples) using population size histories previously inferred using *smc*++ [49,50]. Maps were inferred separately for each chromosome, treating the pseudoautosomal region of the X chromosome as an autosome. For the non-pseudoautosomal region, inferred population sizes were scaled by 0.75 and genotypes from males were combined to create pseudo-diploid samples. In all cases, genotypes were treated as unphased and a mutation rate of 4×10^-9^ mutations per bp per generation was used. This matches the rate implied by the divergence of an ancient wolf sample and is within the confidence interval of the mutation rate estimated from a wolf pedigree [26,51].

Inference using *pyrh*o proceeded in three steps. First, a lookup table of two-locus sampling probabilities conditional on the inferred population size history was generated using *pyrho make_table* using the --*approx* flag with a sample size (-N) set to be 50% larger than the observed number of sequenced chromosomes and with a relative tolerance of 0.1 (--*decimate_rel_tol*). Second, a parameter search was performed using *pyrho hyperparam* to identify values of the window size and block penalty parameters that show the best match to simulated sequences. Parameters that minimize the L2 norm for each population were chosen separately for the autosomes and the X non-pseudoautosomal region. Third, estimates of the per-base recombination rate between SNPs were obtained using *pyrho optimize* with the chosen parameters. The maps inferred for the pseudoautosomal and non-pseudoautosomal regions of the X chromosome were combined and all maps were converted to a centimorgan scale. Finally, each inferred map was rescaled to match the total genetic length of each chromosome inferred by Auton et al.[42] An average genetic map was created based on the average recombination rate between makers inferred from each of the four genetic maps, weighted by the number of chromosomes used in each inference.

### 2.2 Converting previously inferred maps to the canFam4/UU_Cfam_GSD_1.0 assembly

Previous studies generated canine genetic maps relative to the canFam3.1 assembly using linkage disequilibrium and pedigree analysis [42,45]. We converted the published maps to the canFam4 assembly following a stepwise procedure. First, the genetic distances along chromosome 27 and 32 were inverted to account for the orientation differences between the assemblies [41]. Next, physical coordinates from the published maps were converted to canFam4 assembly coordinates using the liftOver tool [52]. Positions that mapped to different chromosomes between the assemblies were discarded and positions were sorted by physical position. The sorted list for each chromosome was then scanned and markers with a smaller genetic map position than the previous marker, or with an implied recombination rate greater than 50 cM/Mb relative to the previous maker, were discarded. The sex-averaged map produced by Campbell et al. was processed using these methods [45].

### 2.3 Correlations of recombination rate between maps

The Spearman correlation between recombination rates from different maps was calculated in 100 bp, 1 kb, 10 kb, 100 kb, 1 Mb, and 5 Mb windows. Only recombination rates on the autosomes were considered. To limit autocorrelation effects over short distances, 100 bp and 1 kb windows were centered every 10 kb.

### 2.4 Recombination hotspot analyses

Recombination hotspots were identified in the inferred genetic maps using the “filtering” method described by Wooldridge and Dumont [53]. Inter-SNP segments with a recombination rate at least 10-times greater than the chromosome average were identified and merged. Segments that include more than two SNPs and are less than 5 kb in size were considered as hotspots. The position of hotspots identified in Auton et al. and of gaps in the canFam3.1 assembly were converted to canFam4 coordinates using liftOver [52]. Overlap between data sets was determined using bedtools intersect [54]. Overlap statistics were limited to the autosomes. Significance of overlaps was determined using bedtools shuffle using the -chrom option to randomly place features on the same chromosome.

### 2.5 Assessment of phasing accuracy

The utility of inferred and converted genetic maps were assessed through an analysis of phasing accuracy. A truth set was constructed using a previously sequenced wolf trio (samples SAMN02921321, SAMN02921320, and SAMN11787846) [51,55]. Illumina sequencing data from each individual was aligned to the UU_Cfam_GSD_1.0 /canFam4 assembly using the same pipeline as the Dog10K Consortium [11]. Genotypes were inferred for each sample at the SNP positions reported in the Dog10K data and the genotype phase of heterozygotes in the child (SAMN02921321) determined following Mendelian segregation rules. Positions with missing data in the trio or a non-Mendelian genotype configuration were discarded. Genotypes for the child were merged with the 1,987 samples included in the Dog10K collection using bcftools [56]. SNPs located in regions identified as segmental duplications were removed. Phased haplotypes were inferred for 1,988 samples (the Dog10K collection plus the wolf child, SAMN02921321) using the *phase_common* utility of SHAPEIT5 (version 5.1.1)[57]. Analysis was limited to biallelic SNPs on chromosome 1 and repeated using the dog average genetic map inferred here, the converted Auton LD and Campbell pedigree maps, and with a constant recombination rate of 1 cM/Mb. Switch errors were tabulated by comparing the allele configuration at consecutive heterozygous SNPs with that inferred from the trio analysis. Since genotype errors may also contribute to apparent phasing errors, switch errors at consecutive positions were defined as paired switch errors and consecutive paired switch errors were labeled as double switches [58]. An estimated phase error rate was calculated based on the number of single and double switches.

## 3. Results

### 3.1 Recombination maps inferred for the canFam4 assembly

Using SNP genotypes from the Dog10K Consortium [11], along with a previously described inference of population size history [50], we inferred recombination maps for four village dog populations using *pyrho* [46,47]. Recombination rates along the non-pseudoautosomal region of the X chromosome were inferred by combining genotypes from female samples with pseudo-diploid genotypes constructed from pairs of males. The inferred maps were constructed from samples of 24-48 chromosomes from each population (Table 1). Internally, *pyrho* estimates recombination in units that depend on the effective population size and mutation rate. Since the absolute scale of these estimates is highly sensitive to modelling assumptions, we rescaled the inferred maps to match the map lengths of each chromosome previously estimated by Auton et al. [42], as done in previous studies [46,59,60]. Additionally, we constructed a consensus recombination map based on the average of the recombination rates inferred for each village dog population weighted by the number of chromosomes used to generate each map. The resulting recombination map is broadly consistent with findings from previous studies in dogs, including a clear increase in inferred rates near the telomeric ends of chromosomes (Figure 1). On the X chromosome, a high recombination rate at the pseudoautosomal region as well as a decreased rate in an extended region that includes the *Xist* locus is seen [61,62].

**Table 1.** Number of samples used from each population. Pseudo-diploid samples constructed using males were included in the analysis of the X chromosome.

| Village<br>Dog<br>Population | Total<br>Samples | Females | Males | Autosomal<br>Maps<br>(chromosomes) | X Map<br>(chromosomes) |
| --- | --- | --- | --- | --- | --- |
| Congo | 16 | 10 | 6 | 32 | 26 |
| Iran | 24 | 9 | 15 | 48 | 32 |
| Kenya | 18 | 7 | 11 | 36 | 24 |
| Liberia | 17 | 12 | 5 | 34 | 28 |

**Figure 1.**
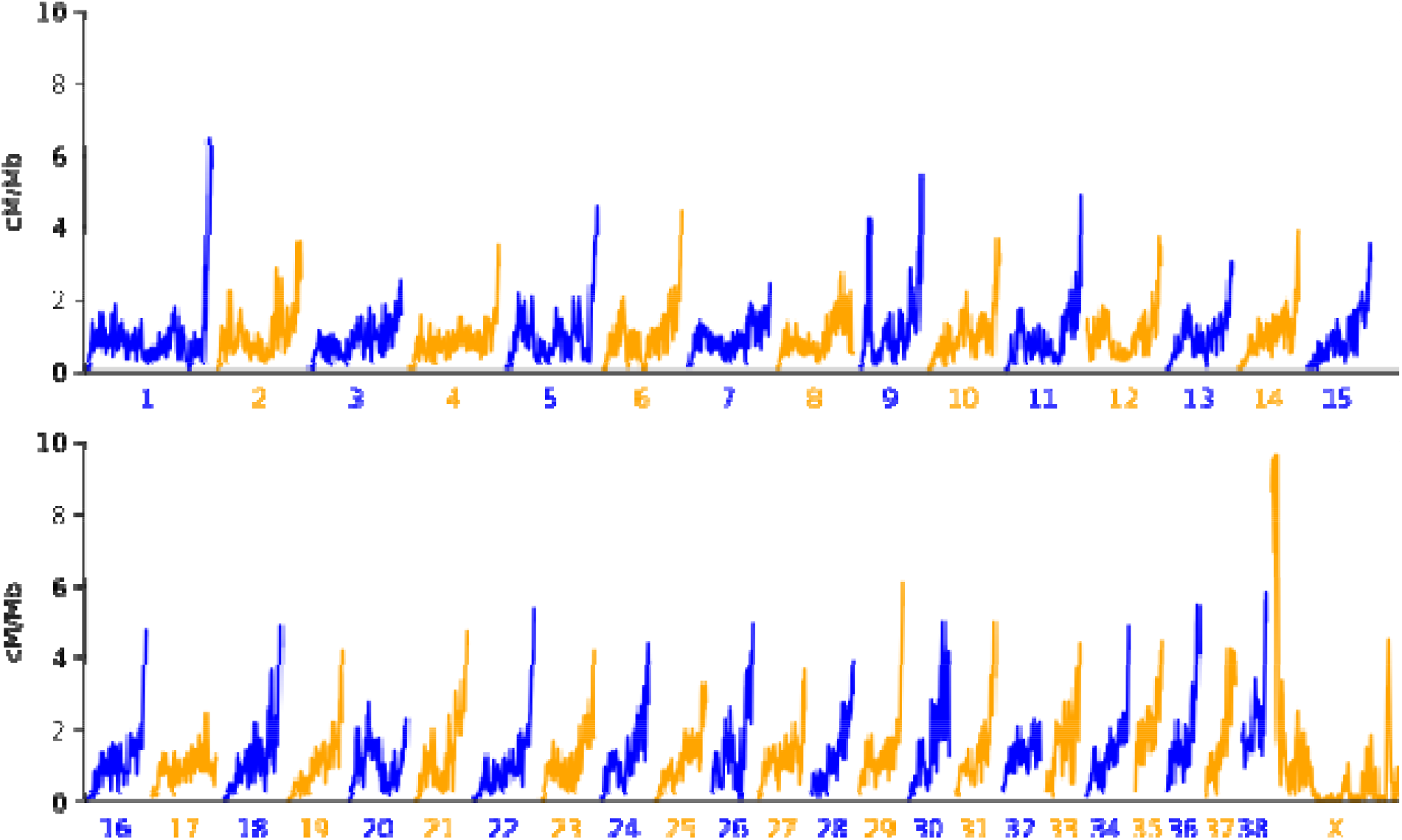
Broad scale recombination rates across the canine genome. The recombination rate from the dog averaged genetic map is plotted in 1 Mb sliding windows across the 38 canine autosomes and the X chromosome. The x-axis scale is constant across plots.

### 3.2 Comparison with previously generated recombination maps

To compare with previously generated canine recombination maps, we converted the recombination rates inferred by Auton et al., based on linkage disequilibrium in village dogs, and by Campbell et al., based on pedigree analysis, to canFam4 coordinates [42,45]. We then calculated the Spearman correlation between pairs of maps at different resolutions (Figure 2). At a broad scale, the rates represented in all maps are highly correlated (Spearman ρ > 0.839 at the 5 Mb scale). At smaller scales, a reduced correlation is found between the LD and pedigree maps. At the 100 bp scale, the correlation between the Campbell pedigree map and the other maps ranges from ρ=0.194-0.222, with the largest correlation found with the dog averaged map. The Auton LD map shows a correlation of ρ=0.611-0.703 with the other LD-based maps, with the largest correlation also found with the dog average map.

**Figure 2.**
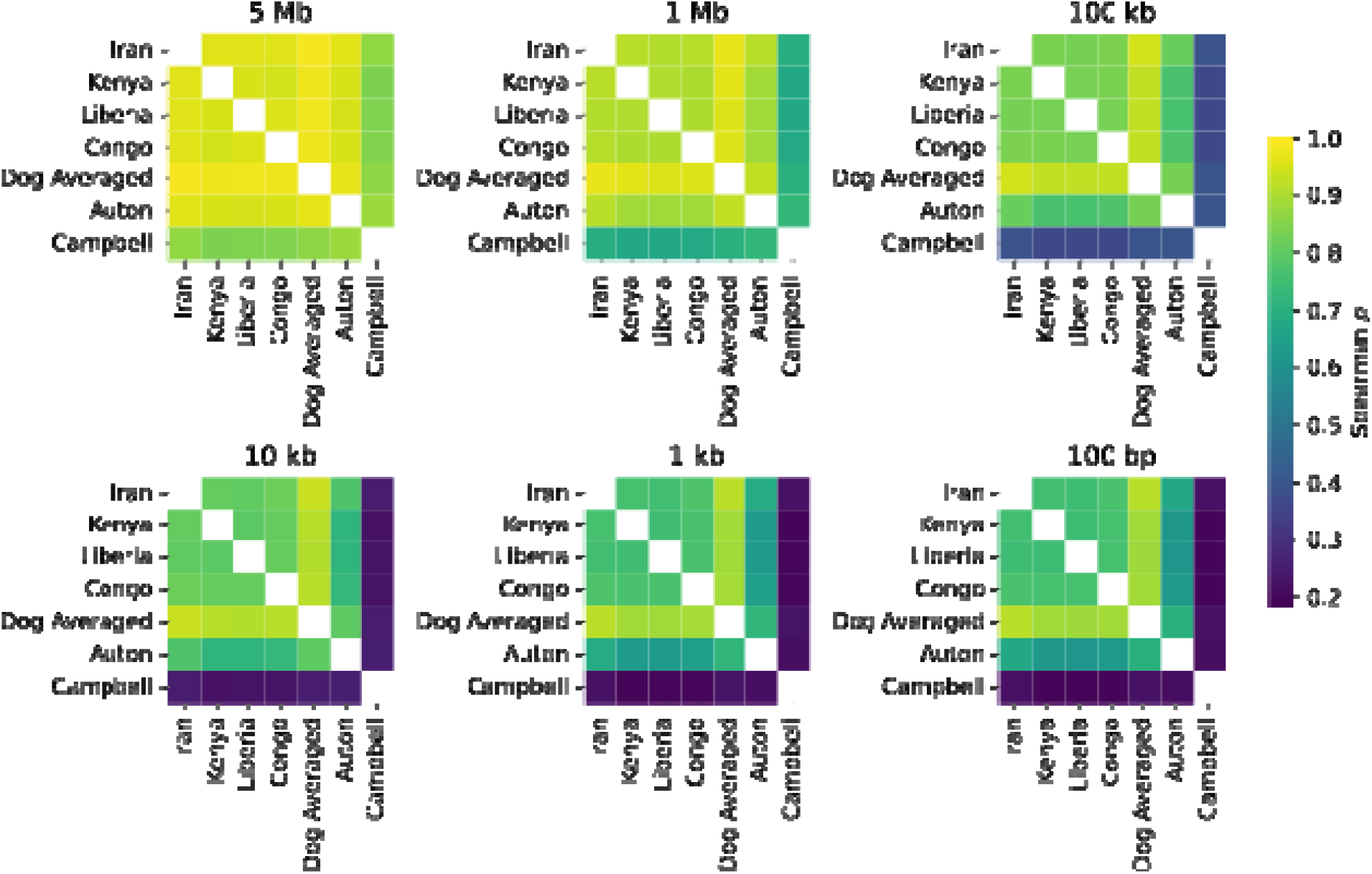
Correlation between genetic maps. Heatmaps depict the Spearman correlation between genetic maps inferred from four village dog populations (Iran, Kenya, Liberia, and Congo), the weighted average dog genetic map, and the converted Auton LD and Campbell pedigree maps. Correlations were calculated in genomic windows 100 bp to 5 Mb in size as indicated.

All analyzed maps show a highly non-uniform distribution of recombination across the canine genome (Figure 3). Consistent with prior findings, the Campbell pedigree map is the most skewed, with 80% of the recombination distance accounted for by only 21.9% of the sequence. This is followed by the Auton LD map (80% of recombination in 26.0% of the sequence) and the four village dog maps we inferred (80% of recombination in 28.9%-31.1% of the sequence). The dog average map is the least skewed, but still shows that only 33.6% of the sequence is required to account for 80% of recombination.

**Figure 3.**
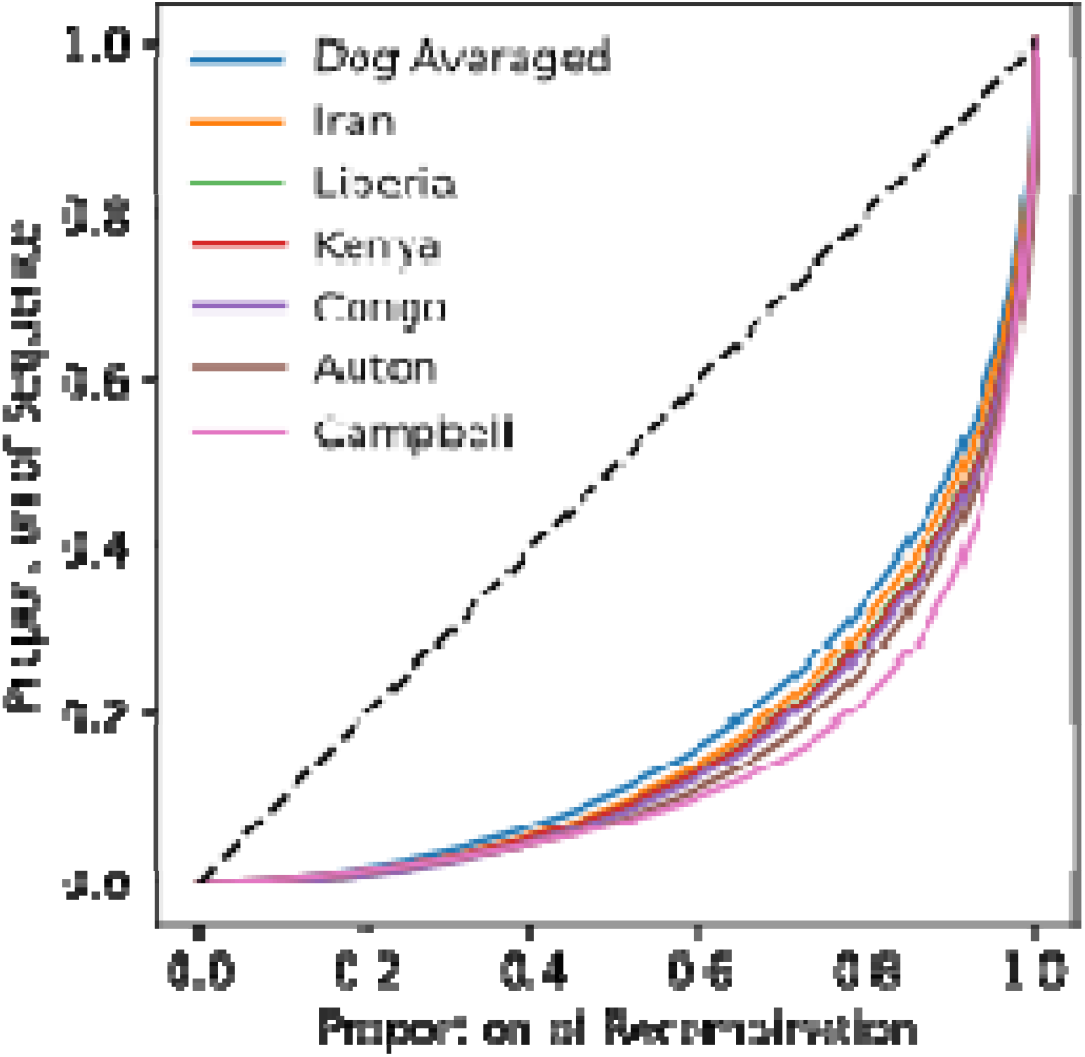
Nonuniform distribution of recombination across the genome and among maps. The proportion of the recombination distance is plotted relative to the proportion of the genome sequence for each analyzed genetic map. The dashed line depicts the pattern expected if recombination is uniform across the genome. Analysis is limited to the autosomes.

Next, we applied a heuristic to identify short segments that have an inferred recombination rate 10-times higher than the chromosome average in each LD map [53]. We then assessed the overlap between the identified hotspots (Figure 4). As expected, the highest overlaps were found between the dog average map and the four maps used to generate it, but 24.7% of hotspots found in the dog average map were also identified in the Auton LD map. This is nearly 11 times greater than the 2.3% of overlaps found by random permutation.

**Figure 4.**
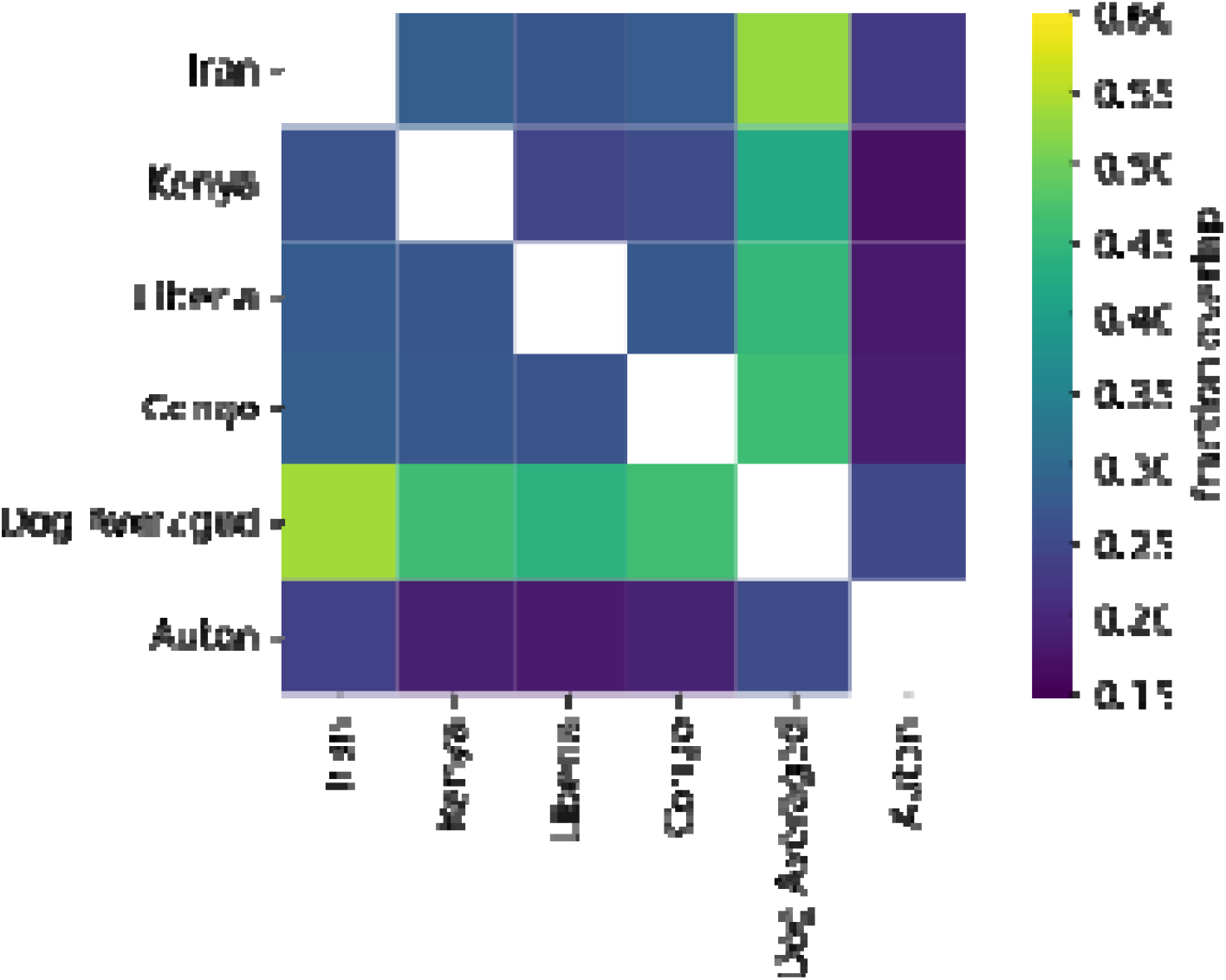
Overlap of hotspots identified using different genetic maps. The heatmap depicts the fraction of hotspots in common between each LD map. Hotspots in the Auton LD map were lifted over to canFam4 coordinates. Values indicate the fraction of hotspots identified for the map listed in each row that overlap with hotspots found from the map listed in each column.

Previous analysis showed that dog recombination hotspots are enriched at regions represented as gaps in the canFam3.1 assembly [13]. The canFam4 assembly, which was built using PacBio sequencing reads, fills in the sequence for most of these gaps [14]. We found that 13.2% of hotspots inferred in the dog average map intersect with regions that are gaps in the canFam3.1 assembly, a 7.8-fold enrichment relative to that expected by chance.

### 3.3 Effect of recombination maps on phasing accuracy

We assessed the effect of different recombination maps on phasing accuracy by phasing genotypes from the offspring of a wolf trio along with the 1,987 samples included in the Dog10K collection. We then compared the inferred genotype phase with the phase determined by Mendelian analysis of the wolf trio. We performed phasing using SHAPEIT5 [57], limited analysis to chr1, and repeated the process using the dog average map, the Auton LD map, the Campbell pedigree map, and a uniform recombination rate. Results consistent with a low phasing error rate were obtained in each analysis (Table 2). Counting every discordancy between consecutive heterozygous positions as a switch error, we estimate the total switch error rate as 2.23% – 2.27%, with the best results obtained using the Auton LD map. Since trio genotyping errors can manifest as phasing errors [58], we additionally identified positions where a single heterozygote is incorrectly phased with respect to flanking positions. Counting double switches as single errors yields estimated phase error rates of 1.62%-1.67%.

**Table 2.** Assessment of phasing accuracy. The number of times the inferred phase for consecutive heterozygotes differed from the phase determined by trio analysis is depicted. A single switch error is an error that is not immediately proceeded or followed by an error. A paired switch error is immediately adjacent to another switch error. Two consecutive paired switch errors are considered to be a double switch error. The estimated phase error rate is derived from the number of single and double switches.

| Genetic<br>map | Mendelian-<br>phased<br>heterozygotes | Switch<br>errors | Switch<br>error rate | Single<br>switches<br>+ double<br>switches | Phase error<br>rate |
| --- | --- | --- | --- | --- | --- |
| Auton | 98765 | 2205 | 0.0223 | 1602 | 0.0162 |
| Dog<br>Average | 98765 | 2233 | 0.0226 | 1650 | 0.0167 |
| Campbell | 98765 | 2241 | 0.0227 | 1640 | 0.0166 |
| Constant<br>Rate | 98765 | 2243 | 0.0227 | 1649 | 0.0167 |

## 4. Discussion

Recent advances have enabled the inference of fine-scale recombination maps from populations with a changing effective population size [46]. We combined these advances with a recently generated catalog of canine variation identified using Illumina short-read sequencing. Using *pyrho*, we inferred new recombination maps based on linkage disequilibrium pattens found in 16-24 village dogs sampled from four populations. We also systematically converted two previously published maps to the coordinates represented by the new canFam4 assembly. All maps are strongly correlated at large scales, while the pedigree-based map shows reduced correlation at finer scales.

As in other species that lack a functional copy of *Prdm9*, recombination in dogs appears concentrated in regions near the beginning of genes and in regions with a high GC content [36,42]. Previous studies of recombination in dogs relied upon the canFam3.1 reference, which contains over 19,000 gaps where the underlying sequence is unknown. Assemblies constructed using long-read sequencing technologies filled in most of these gaps, revealing that they contained sequences with an extremely high GC content [13]. Using the fine-scale genetic map directly inferred using the canFam4 assembly, we confirm that canine recombination hotspots are enriched at these regions, consistent with the contribution of GC-biased gene conversion at recombination hotspots to the evolution of canine genome structure [13,36,42].

To assess the utility of the new and previously published maps we performed an assessment of haplotype phasing accuracy. A phase error rate of ∼2% was found when using the samples from the Dog10K collection as a reference panel, with choice of genetic map having little effect on the result. This is consistent with the notion that reference panel size is a major driver of phasing accuracy [63]. The Dog10K collection includes breed dogs, village dogs, and Eurasian wolves while the wolf trio we used as a benchmark was sampled from North America [51,55]. Thus, our analysis represents results that may be expected when using a sample from a population not included in the reference panel. Higher accuracy may be obtained using samples that are more closely related to the panel.

It may be surprising that the Auton LD map showed the lowest phase error rate in our analysis. Several factors may contribute to this. We caution that these findings are based on the use of a single sample as a truth set and that the differences in estimated error rate are very modest. Although *pyrho*, which considers changes in population size history, is expected to offer improved inference relative to *LDhat*, sample size may also play an important role [46]. Our analysis inferred maps separately from four populations, each with 16-24 samples. In contrast, Auton et al. created a single map based on 51 village dogs collected from diverse locations [42]. The benefits of using a sample twice as large while ignoring population structure and demographic history may outweigh the application of a more sophisticated method to a smaller sample. Additionally, for over 15 years *LDhat* has been the most widely used method for inferring fine-scale recombination maps [43,44]. As result, the parameters of phasing algorithms may have been tuned to maximize performance based on the characteristics of maps generated by *LDhat*.

The expected recombination fraction between genetic markers is an important parameter in many analyses. The canine genomics community has largely transitioned to the use of new long-read genome assemblies, and updating key resources is critical for adoption of improved references. The genetic maps described here, including previously published maps uniformly transitioned to the canFam4 assembly, will be useful for future studies including scans of selection, measures of haplotype sharing, the inference of population sizes, and the imputation of missing genotypes.

## Author Contributions

J.M.K. designed the study, performed analysis, and wrote the manuscript.

## Data Availability Statement

Genetic maps described in this study are available from the Zenodo data archive under accession DOI 10.5281/zenodo.17095604.

## Acknowledgments

I thank Matthew S. Blacksmith and Emily C. Koch for constructive comments on a draft of this manuscript.

